# Viral non-coding RNA structure annotation and API-based data retrieval with Rfam and R2DT

**DOI:** 10.64898/2026.05.10.724034

**Authors:** Philippa Muston, Sandra Triebel, Eric Nawrocki, Nancy Ontiveros-Palacios, Isaac Jandalala, Blake Sweeney, Alex Bateman, Manja Marz, Anton I. Petrov, Pedro Madrigal

**Affiliations:** European Molecular Biology Laboratory, European Bioinformatics Institute, EMBL-EBI, Hinxton, United Kingdom; Friedrich Schiller University Jena, Germany; National Library of Medicine, National Institutes of Health, Bethesda, United States; RNera Bio Ltd, Cambridge, United Kingdom; Riboscope Ltd, Cambridge, United Kingdom

**Keywords:** RNA families, Sequence database, API, Secondary structure, Rfam, R2DT, Infernal, Viruses, ncRNA annotation

## Abstract

Rfam is a comprehensive database of non-coding RNA (ncRNA) families providing curated sequence alignments, consensus secondary structures, and covariance models for thousands of RNA families. The database is essential for identifying structured non-coding RNAs in newly sequenced genomes and understanding RNA structure-function relationships. Here we present computational protocols for automated ncRNA annotation of viral genomes, and for programmatic interaction with Rfam through its RESTful API. We showcase genome-wide RNA structure visualization from a genome sequence and from a multiple sequence alignment by generating comprehensive 2D structure diagrams using newly developed features in R2DT. We also present practical examples for retrieving family metadata, downloading alignments, accessing secondary structures, and searching user sequences from the Rfam API. These methods enable researchers in virology and RNA biology to integrate Rfam data into custom bioinformatics pipelines, comparative analyses, and machine learning workflows.

## 1 Introduction to the Rfam database

The Rfam database is a central repository of non-coding RNA (ncRNA) families, which started in 2002 and has since been continuously maintained [1,2]. Each family in Rfam is represented by a multiple sequence alignment (MSA), also called SEED, and a covariance model (CM). The SEED alignment contains aligned homologous RNA sequences that share a consensus secondary structure [1]. The family includes complementary information such as references for the alignment and secondary structure source, ontologies, and a general description linked to a Wikipedia entry [3]. The CM is trained on the SEED alignment and built using the Infernal software [4]. The FULL hits are a set of matches found using the CM for each family, obtained using Rfamseq, a comprehensive collection of whole genomes. For Rfam 15.0, the reference genome collection includes 26,106 non-viral and 13,552 viral genomes [2]. The searches return a list of hits, or putative homologs, ranked by bit scores derived from the CMs. The FULL hit set is defined after the selection of a threshold that discriminates between *bona fide* homologs to the seed sequences and the background distribution of false positive hits.

The latest version of Rfam (15.1, January 2026) has 4,227 families. Rfam families are based on MSAs annotated with consensus secondary structures that indicate base pairing. The accuracy of consensus secondary structure is important as it guides the SEED alignment and informs the Infernal CM. Secondary structures in Rfam have been determined by biochemical studies or bioinformatic prediction. Whenever 3D information is available, Rfam also updates family structures with the available 3D information from PDB [5]. Families reviewed with 3D information can be accessed at https://rfam.org/3d. ncRNA alignments provided by Rfam are a valuable resource for annotating and exploring the structure and distribution of ncRNAs.

Viral RNA genomes, ranging from 3.4 kb to 41 kb, contain highly conserved RNA structures that serve vital roles in exonuclease resistance, genome diversification, and multiple stages of the viral replication cycle. More recent viral RNA families in Rfam include *Coronaviridae* and *Flaviviridae* families in Rfam 14.3 [6], hepatitis C virus in Rfam 15.0 [2] and *Pestivirus* families in Rfam 15.1 [7]. Recent viral Rfam families can be accessed at https://rfam.org/viruses. The project is done in collaboration with expert virologists from the European Virus Bioinformatics Centre (EVBC). In previous years, the EVBC team has produced curated, representative and non-redundant genome-wide alignments for several viral families. The review and update of viral families is an ongoing effort aimed at providing a comprehensive dataset for virologists, enabling the annotation of viral genomes with conserved RNA structures using Infernal software [4] and Rfam covariance models [8].

In this chapter, we present a protocol to visualise Rfam structures in the context of the viral genome using the R2DT framework for predicting and visualising secondary structures using templates [9,10]. R2DT version 2.2 and later accept either a Stockholm alignment file or a raw genome sequence as input and generates diagrams showing the location of the Rfam hits within the genome and their secondary structures. Improved understanding of RNA structure in viral pathogens affecting animals, plants and humans could lead to improved disease control, prevention and preparation for future pandemics, and the development of new vaccines to fight viruses.

Rfam is commonly used as a data source for training AI models to predict functions, structure and interactions of ncRNAs [11]. To showcase data retrieval, we also present a protocol for programmatic access to data using the Rfam REST API. Browsing and searching families, genomes, and sequences via the website, FTP, or public MySQL database have already been covered in a 2018 Rfam protocol paper [12]. By covering here Rfam data access via RESTful API endpoints and programmatic HTTP access with JSON/XML data retrieval, we provide scalable, on-demand data access that supports cloud-native applications, automated annotation pipelines, and real-time integration with machine learning frameworks. Researchers can efficiently query and retrieve ncRNA family data programmatically, eliminating the need for bulk downloads or local database maintenance.

## 2 Viral annotation of genomes and multiple sequence alignments

### 2.1 ncRNA annotation of viruses using Rfam/Infernal

Annotation of structured RNAs in genomes and other sequence datasets provides insight into regulatory and functional elements encoded in the organisms represented in those datasets. The Infernal software package includes programs for annotating structural RNAs in sequence datasets from all domains of life using covariance models (CMs) that include both sequence and secondary structure conservation information [8]. Here we focus on how to use Rfam models with Infernal to annotate viral sequence datasets.

#### 2.1.1 Extract viral Rfam models

The first step is to extract the subset of Rfam models that include viral sequences. One could use all Rfam models but that would require more time and computational resources and would introduce potential false positives and/or Rfam hits to contaminated sequences. Instead, it is recommended to download the Rfam CM file and the `viruses.clanin` file that includes information on virus ‘clans’ (groups of related Rfam families) and use the information in the rfam-taxonomy GitHub repository to extract the CMs specific to viruses and create a `viruses.cm` file:

~~~
$curl -L -o Rfam.cm.gz https://ftp.ebi.ac.uk/pub/databases/Rfam/15.1/Rfam.cm.gz
$curl -L -o viruses.clanin
 https://raw.githubusercontent.com/Rfam/rfam-taxonomy/master/domains/viruses.clanin
$curl https://raw.githubusercontent.com/Rfam/rfam-taxonomy/master/domains/viruses.csv | cut
 -f 1,1 -d ‘,’ | tail -n +2 | cmfetch -o viruses.cm -f Rfam.cm.gz –
~~~

#### 2.1.2 Run cmscan to annotate structural RNAs

As an example dataset, download all complete genomes of RNA viruses (*Riboviria*) in the RefSeq database using the NCBI Datasets tool [13]:

~~~
$datasets download virus genome taxon Riboviria --refseq --complete-only --filename
 riboviria_refseq_complete.zip; unzip riboviria_refseq_complete.zip; mv
 ncbi_dataset/data/genomic.fna viruses.fa
~~~

To annotate matches to Rfam families in `viruses.fa` in FASTA format, run Infernal’s `cmpress` tool to prepare the CM file and then run the `cmscan`tool:

~~~
$cmpress viruses.cm
$cmscan --nohmmonly --rfam --cut_ga --fmt 2 --oclan --oskip --clanin
 viruses.clanin -o viruses.cmscan.out --tblout viruses.cmscan.tblout viruses.cm
 viruses.fa
~~~

Infernal (`cmfetch, cmpress, cmscan`) and NCBI datasets installation instructions can be found at http://eddylab.org/infernal/ and https://www.ncbi.nlm.nih.gov/datasets/docs/command-line-start/, respectively.

#### 2.1.3 Analyse the output

All matches to models with scores above the manually curated Rfam family-specific GA score thresholds (`--cut_ga`) will be output in tabular form to `viruses.cmscan.tblout` and in more verbose form including alignments to `viruses.cmscan.out`, as described in the Infernal user’s guide (http://eddylab.org/infernal/Userguide.pdf). For the `viruses.fa` file returned by Datasets at the time of writing, which has 9146 sequences, there are 1311 hits to 164 families in the `viruses.cmscan.tblout` file. The matches to each sequence will be sorted by E-value, with all hits to the first sequence followed by all hits to the second sequence, and so forth. To generate output on a per-family basis (all hits to each family grouped together), use the `cmsearch` program with the same flags as above for `cmscan` except omit `--fmt 2, --oclan, --oskip` or `--clanin viruses.clanin`. With `cmsearch`, lower scoring overlapping hits for models in the same Rfam clan (groups of homologous models, e.g. CL00117 which includes seven coronavirus 3’ UTR families) are not removed, but such hits are removed by `cmscan`. An example hit alignment for the Flavivirus DB element family (RF00525) from `viruses.cmscan.out` is shown below:

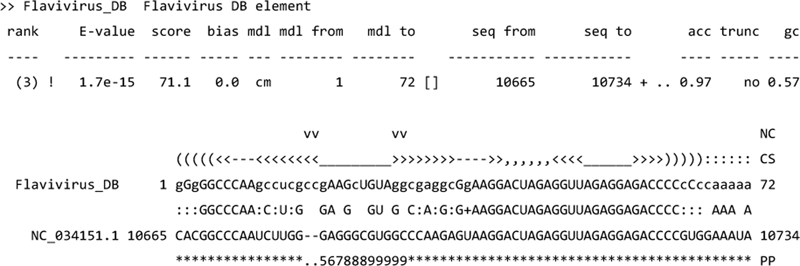

The example shows a match in the `viruses.fa` sequence dataset to RF00525 (Flavirus_DB) in the NC_034151.1 genome of the T’Ho virus. For explanation of the alignment format see the tutorial section of http://eddylab.org/infernal/Userguide.pdf.

### 2.2 RNA annotation from a genome sequence

In this section, we switch focus to individual viral genomes and describe a workflow that combines Infernal searches against Rfam covariance models with R2DT visualisation to generate a genome-wide two-dimensional (2D) RNA structure map. This approach enables consistent, template-based visualisation of RNA secondary structures detected across the viral genome. The end-to-end analysis can be performed using R2DT version 2.2 or higher from a genome sequence in FASTA format.

#### 2.2.1 Prepare the input

This example uses the SARS-CoV-2 genome with accession OX309346.1, which is included in the R2DT repository at examples/viral/coronavirus.fasta^1^.

#### 2.2.2 Run viral-annotate in R2DT

R2DT uses Infernal’s `cmscan` tool to search viral genomes against the Rfam covariance model library, then generates secondary structure diagrams for each identified RNA family.

`$r2dt.py viral-annotate examples/viral/coronavirus.fasta output/coronavirus/`

#### 2.2.3 Examine the output

The`output/coronavirus/` directory contains:

~~~
output/coronavirus/
⊢ cmscan.tblout # Tabular cmscan results
⊢ cmscan.out # Full cmscan output
└ rfam/ # Individual RNA diagrams
 ⊢ RF03120_26-299.colored.svg
 ⊢ RF00507_13469-13546.colored.svg
 └ RF03125_29536-29870.colored.svg
~~~

Each SVG file shows the secondary structure of one RNA element. An example for RF03120 is shown in Figure 1.

**Figure 1.**
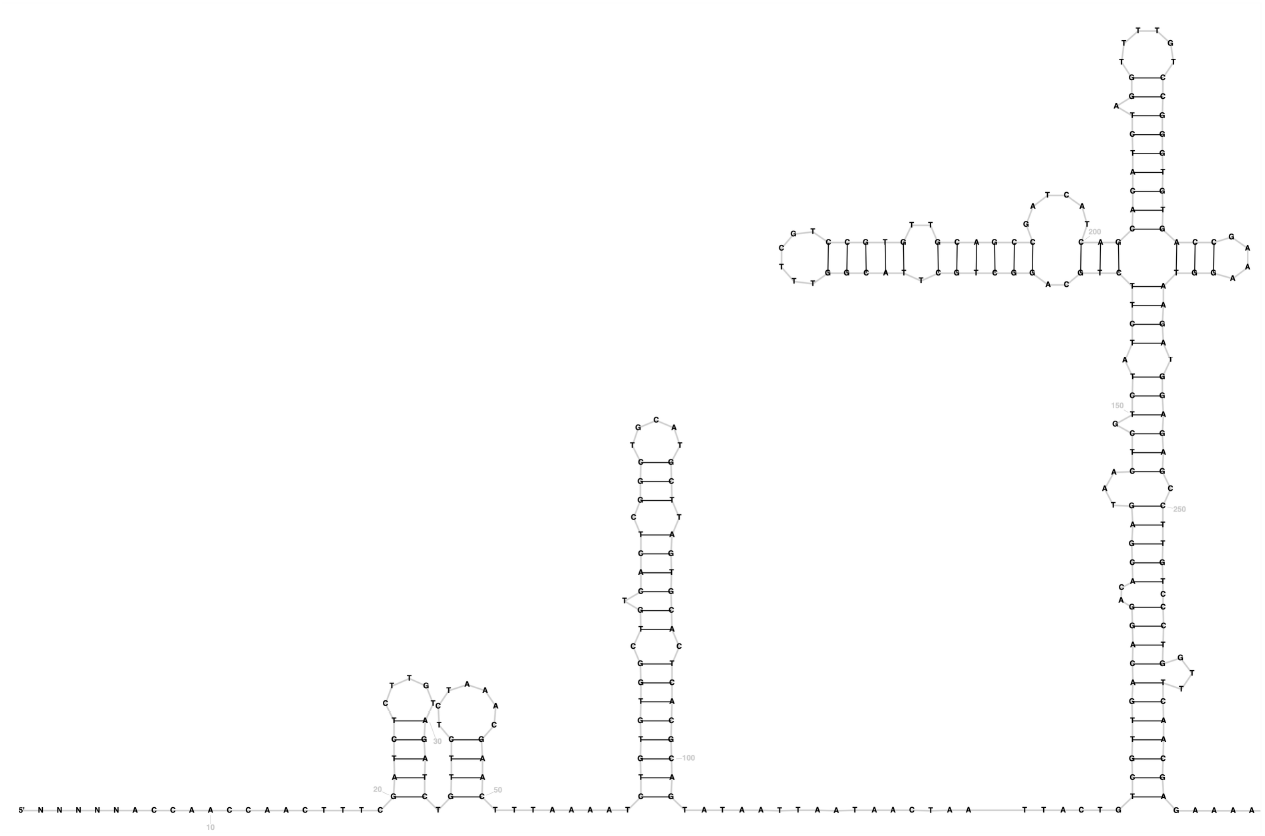
RF03120 (Sarbecovirus-5UTR): The 5′ untranslated region containing stem-loops SL1-SL5, 5′ UTR of SARS-CoV-2 (nucleotides 26-299).

#### 2.2.4 Create a combined stitched diagram

Finally, combine all diagrams into a single panoramic view:

~~~
$r2dt.py stitch output/coronavirus/rfam/*.colored.svg **\**
 -o coronavirus-stitched.svg --sort **\**
 --captions “5′ UTR” --captions “FSE” --captions “3′ UTR”
~~~

The `--sort` flag arranges panels by genomic coordinates (extracted from filenames), and `-- captions` adds labels below each panel. Each Infernal hit is rendered using its corresponding Rfam reference structure, ensuring visual consistency across all instances of the same RNA family. The resulting stitched diagram shows all three RNA structures in genomic order in Fig. 2.

**Figure 2.**
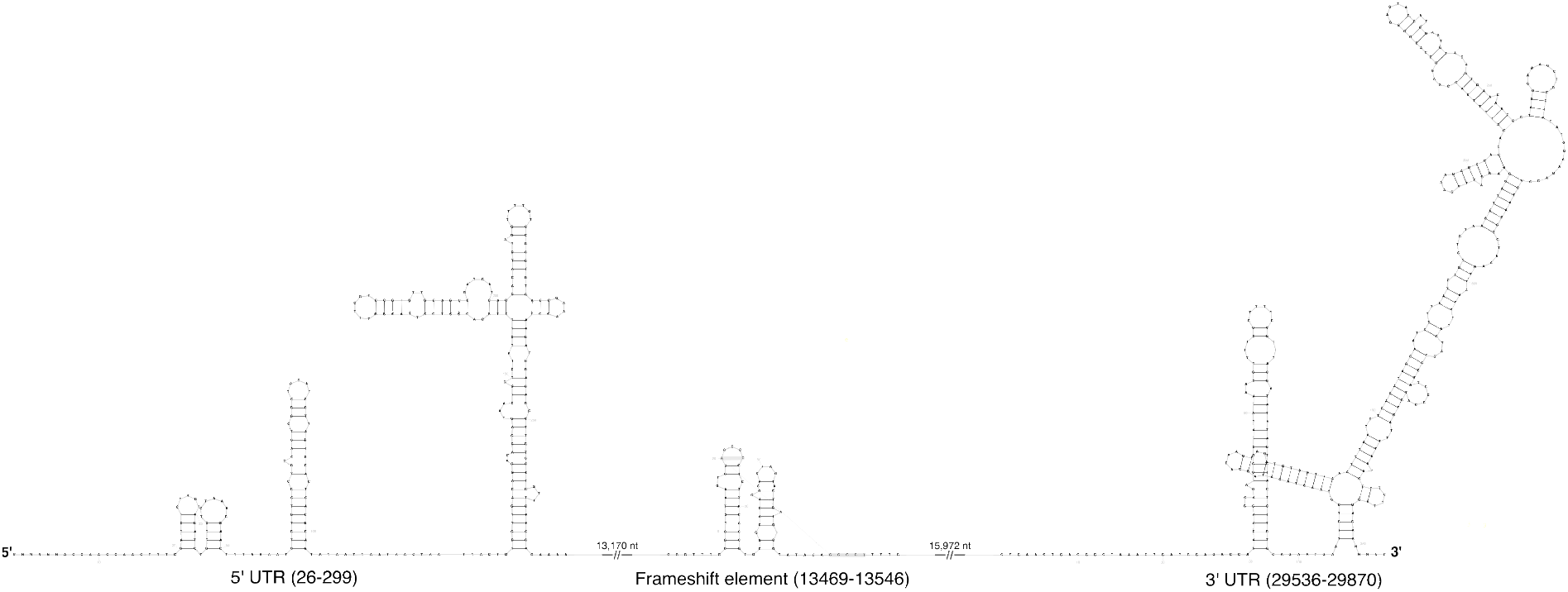
Combined view of SARS-CoV-2 RNA structures: 5′ UTR, FSE (frameshift element), and 3′ UTR.

More examples of annotated viral sequences can be found at https://docs.r2dt.bio/en/latest/viral-genomes.html.

### 2.3 Viral annotation from an MSA

RNA secondary structure prediction based on a single sequence is inherently limited. Thermodynamic folding algorithms typically return one or a small number of energetically favourable structures, but these predictions lack evolutionary context and are often ambiguous, particularly for long RNAs. In contrast, comparative approaches based on multiple sequence alignments (MSAs) exploit evolutionary information to identify structurally conserved elements that are more likely to be biologically relevant [14].

The key advantage of using multiple sequences lies in the detection of structural conservation despite sequence variation. Functional RNA secondary structures tend to be preserved over evolutionary time, even as individual nucleotides mutate. Compensatory and semi-compensatory substitutions that maintain base pairing provide strong evidence for conserved helices and cannot be identified from a single sequence alone. Thus, MSAs enable the distinction between incidental folding and evolutionarily selected RNA structures.

Our understanding of RNA secondary structure in viruses remains incomplete, particularly within protein-coding regions. Historically, structured RNAs have been studied mainly in untranslated regions (UTRs), where their regulatory roles are well established [15,16].

However, growing evidence suggests that structured elements are also widespread within coding regions, where they may influence replication, translation, genome packaging, or host interactions [17-19].

Viral RNA genomes pose specific challenges for secondary structure analysis. They are highly variable at the sequence level and constrained by overlapping coding requirements. These features make reliable single-sequence structure prediction especially difficult. Comparative approaches that leverage homologous viral genomes are therefore essential to uncover conserved RNA structures that are not apparent from sequence analysis alone.

As an alternative to starting from a genome sequence, we now describe a protocol to generate an MSA for viruses that are not yet fully integrated into Rfam, extract structural regions, create covariance models and R2DT templates, and generate 2D diagrams.

#### 2.3.1 Generating multiple sequence alignments from genomes

Triebel and colleagues demonstrated that it is feasible to generate full-genome multiple sequence alignments annotated with RNA secondary structure for viral clades, including hepatitis C virus (*Hepacivirus hominis*, HCV, species level) [7] and *Pestivirus* (genus level) [20]. Their workflow integrates sequence alignment and RNA structure prediction in a scalable and evolution-aware manner, enabling the systematic identification of conserved structural elements across entire viral genomes. The workflow consists of the following steps (Fig. 3):

**Figure 3.**
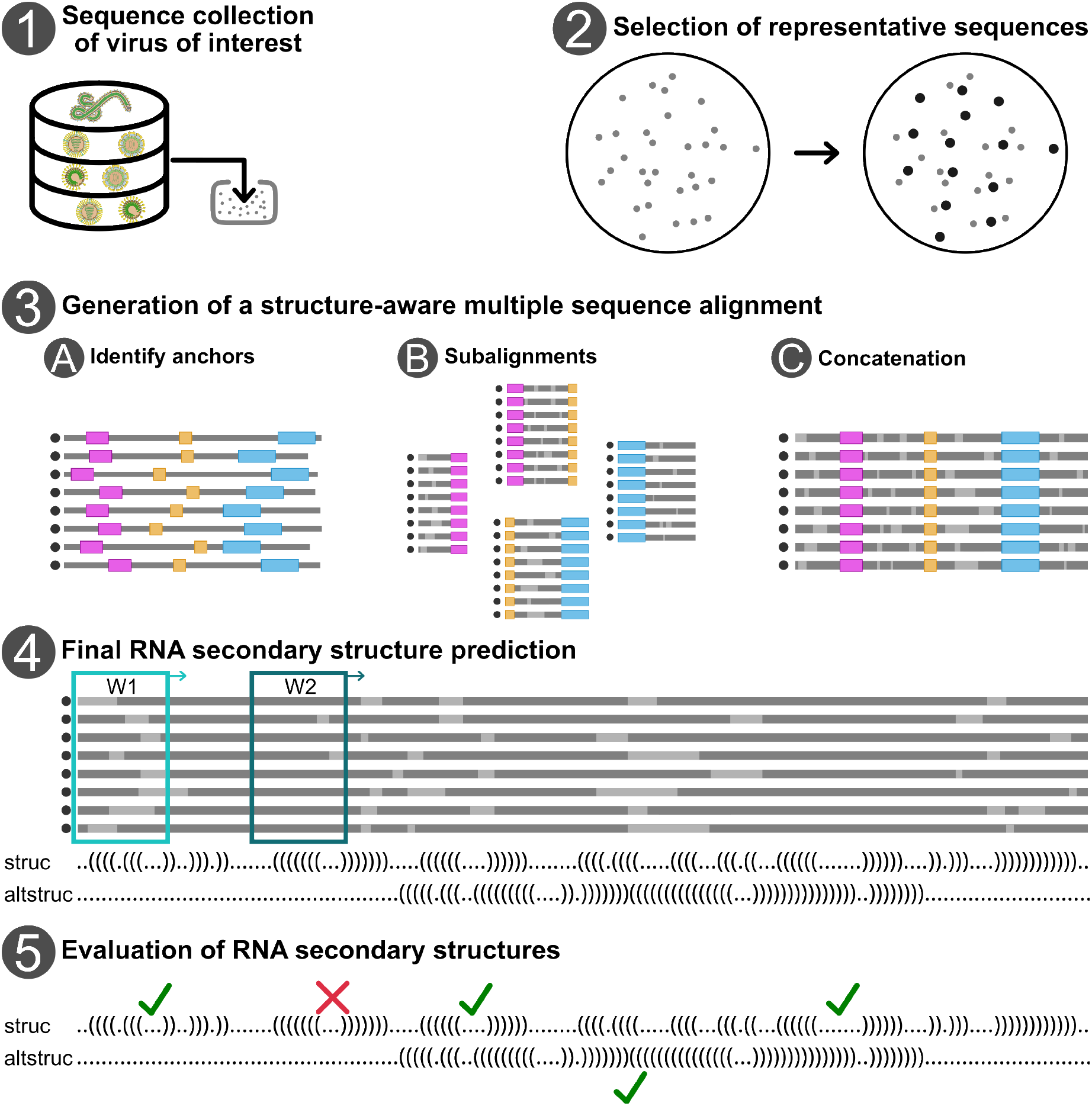
Computational workflow to generate representative multiple sequence alignments with RNA secondary structure annotation for a collection of viral genome sequences.

1. **Sequence collection of the virus of interest** Download genome sequences (FASTA file) and annotation (GFF file) of the virus or viral clade of interest from public databases, e.g. NCBI Virus (https://www.ncbi.nlm.nih.gov/labs/virus/vssi/#/) or BV-BRC (https://www.bv-brc.org/). We recommend searching for the taxonomic name of the virus and using two filters: (1) for genome/nucleotide completeness, to obtain complete genomes (including UTRs and protein-coding regions) or coding-complete genomes (protein-coding regions only); (2) to remove lab strains (lab passaged) because these sequences might have significant differences that do not naturally occur ^2^.

~~~
$datasets download virus genome taxon ‘Hepacivirus hominis’ --complete-only --filename
hepacivirus_hominis.zip
~~~ We will use HCV sequences in FASTA format as an example throughout the rest of this section.
2. **Selection of representative sequences** Cluster the dataset to reduce the number of genomes for subsequent steps by selecting representative sequences, for example using ViralClust, a modular Nextflow pipeline for bias-aware selection of representatives from large viral genome datasets [21]. ViralClust incorporates 5 different clustering algorithms (`CD-HIT-EST, SUMACLUST, VSEARCH, MMSeqs2`, and `HDBSCAN`) enabling comprehensive comparison of the results. This helps to remove redundancy and over-representation while preserving the sequence space adequately. The Nextflow workflow and documentation for ViralClust is available at https://github.com/rnajena/viralclust.

~~~
$nextflow run viralclust.nf --fasta hepacivirus_hominis.fasta --output
viralclust_hepacivirus_hominis/ --eval –ncbi
~~~
3. **Generation of a structure-aware multiple sequence alignment**
  a. Identify conserved regions (called ‘anchors’) present in all genomes, e.g. using AnchoRNA [22]. Anchors are calculated on amino-acid level sequences because of the higher sequence similarity. Parameters regarding the minimum length of an anchor and a minimum BLOSUM62 score between each anchor can be specified in the `config.json` file. The genome sequences are then cut at the anchors to divide the alignment construction into smaller tasks.

~~~
$# calculate anchors
$anchorna go -c conf.json hepacivirus_hominis.gff anchorna_hepacivirus_hominis.gff
$# extract region between anchors, e.g. anchors A0 and A1
$anchorna cutout -o anchorna_hepacivirus_hominis_A0_A1.fasta -m seqoffset
hepacivirus_hominis.fasta anchorna_hepacivirus_hominis.gff A0 A1
~~~
  b. Generate subalignments between anchors. For a sequence and structure aware alignment construction, LocARNA [23] can be utilized. Thus, we obtain an alignment and predicted local RNA secondary structures as a result. We recommend including anchors at the start and end of each alignment. The conserved regions tend to improve the performance of alignment tools.

~~~
$mlocarna --stockholm --consensus-structure alifold --keep-sequence-order --tgtdir
hepacivirus_hominis_A0_A1_mlocarna/ anchorna_hepacivirus_hominis_A0_A1.fasta
~~~
  c. Concatenate subalignments to produce a full-genome multiple sequence alignment. Anchor regions at the start and end of each alignment that have been aligned twice are only included once in the final alignment. Overlapping structures are preserved for later analysis.

~~~
$python anchorna-alimerge -o hepacivirus_hominis_merged.stk <input1.stk> <input2.stk>
<input3.stk> …
~~~
4. **Final RNA secondary structure prediction** Infer consensus RNA secondary structures from the full genome alignment by integrating structural information from all subregions for: (1) global structures - by first applying complexity and coverage filters, followed by a double sliding window approach (W1 and W2, see Fig. 3) for structure prediction using LRIscan [24], (2) local structures – by using a single sliding window (W1) that allows a maximum base pair span of *L* using the ViennaRNA package (e.g., RNALalifold) [25]. `$RNALalifold -p --noLP -f S hepacivirus_hominis_merged.stk` For a coloured output (number of base pair types), add the following parameters: `$--color –aln` To allow colouring based on the number or percentage of sequences folding into the predicted structure, add the following parameter: `$--color--threshold=<number>` Values between 0 and 1 are treated as frequencies among aligned sequences. All other values are truncated to integers and used as absolute counts.
5. **Evaluation of RNA secondary structures** Examine and evaluate predicted RNA structures, especially in regions with overlapping structures and other functional elements, taking into account previous studies, covariation, e.g., using R-scape [26], and conservation beyond the requirements of the coding sequence.

The result is a full-genome multiple sequence alignment annotated with RNA secondary structure (`hepacivirus_hominis_alignment.stk`), whose visualisation is shown for HCV in Fig. 4. More details can be found in the study of Triebel *et al*. [7], where the resulting alignment with RNA secondary structures was submitted as candidate families to the Rfam database, such as the family RF00061, https://rfam.org/family/RF00061 ^3^.

**Figure 4.**
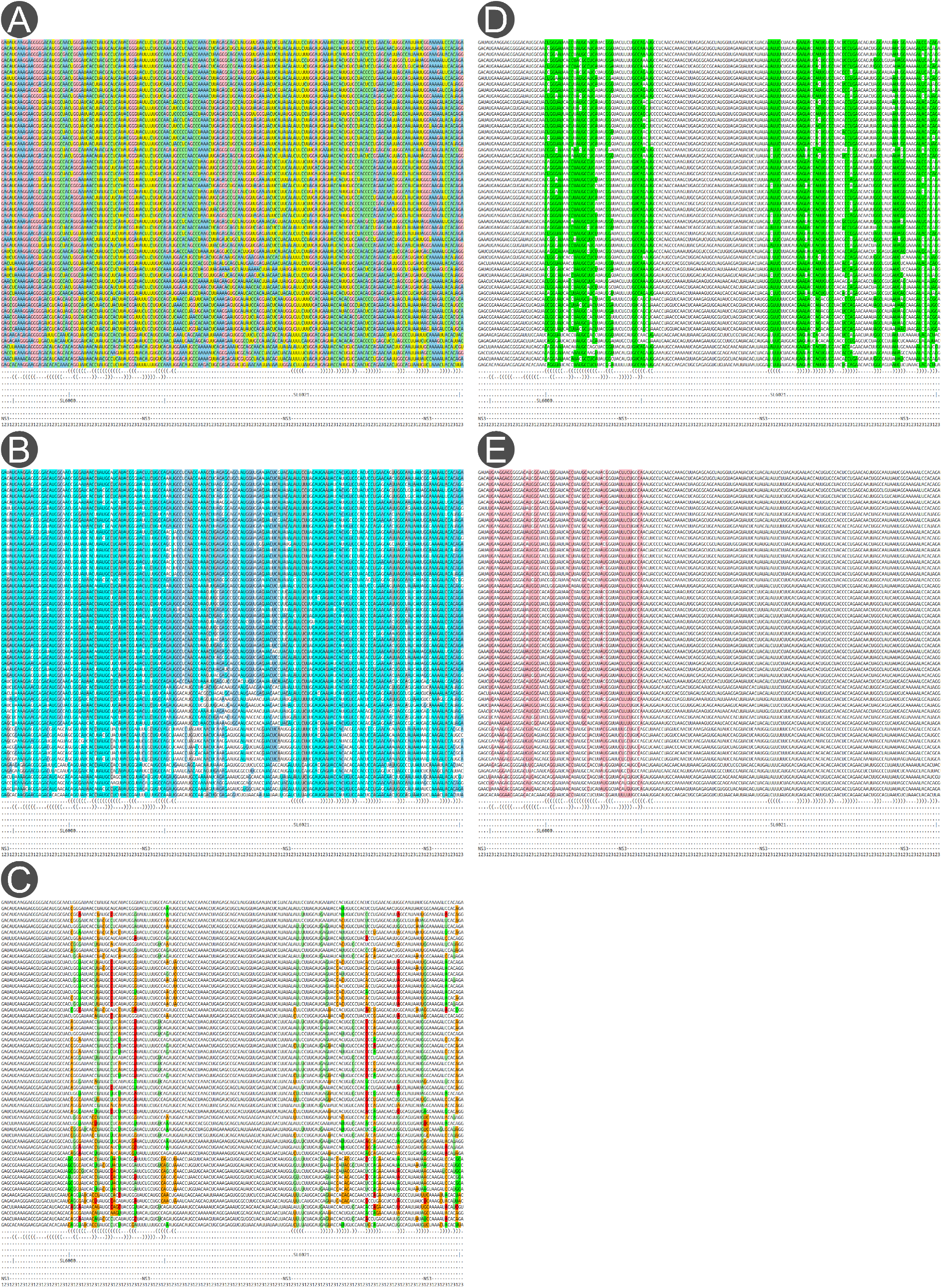
Nucleotide alignment of HCV visualised using Emacs in RALEE mode displaying **(A)** nucleotide identity, **(B)** sequence conservation, **(C)** compensatory mutations, **(D)** consensus RNA secondary structure, and **(E)** alternative RNA secondary structure.

#### 2.3.2 Visualisation of RNA structures in annotated alignments

In addition to sources of data for Rfam, such structure-annotated alignments provide a rich resource for downstream analyses. When used as input for R2DT, they enable visualisation of conserved RNA secondary structures in two-dimensional layouts for specific structures or genome-wide. This facilitates intuitive inspection of structural elements across the viral genome and supports comparative and functional interpretation.

The R2DT software automatically generates RNA secondary structure diagrams in consistent, reproducible and recognisable layouts using a library of templates representing a wide range of RNAs [9-10]. In v2.2 and later versions, R2DT automatically identifies and visualises non-coding RNA structures in viral genomes. R2DT uses the Rfam covariance model database, finding regulatory elements like UTRs, frameshift signals, IRES, and packaging sequences. For Stockholm alignments used as input, each RNA element must have its secondary structure boundaries specified in the `#=GC regionID` line, with individual elements separated by pipe (`|`) characters. For example:

~~~
#=GC SS_cons ............((((((.))))))..
#=GC structureID ............|.......SLI....|..
#=GC regionID |...................5’UTR.....
~~~

This excerpt from the Stockholm file indicates that the SLI hairpin spans positions 14–28 in the alignment, and forms part of the broader 5′ UTR element. The closing pipe character is omitted here because the full 5′ UTR annotation is much longer. R2DT parses these annotation lines automatically, using `structureID` entries as labels for individual structural elements and `regionID` entries for higher-level region names.

For the Hepacivirus Stockholm alignment obtained before (see Fig. 5):

**Figure 5.**
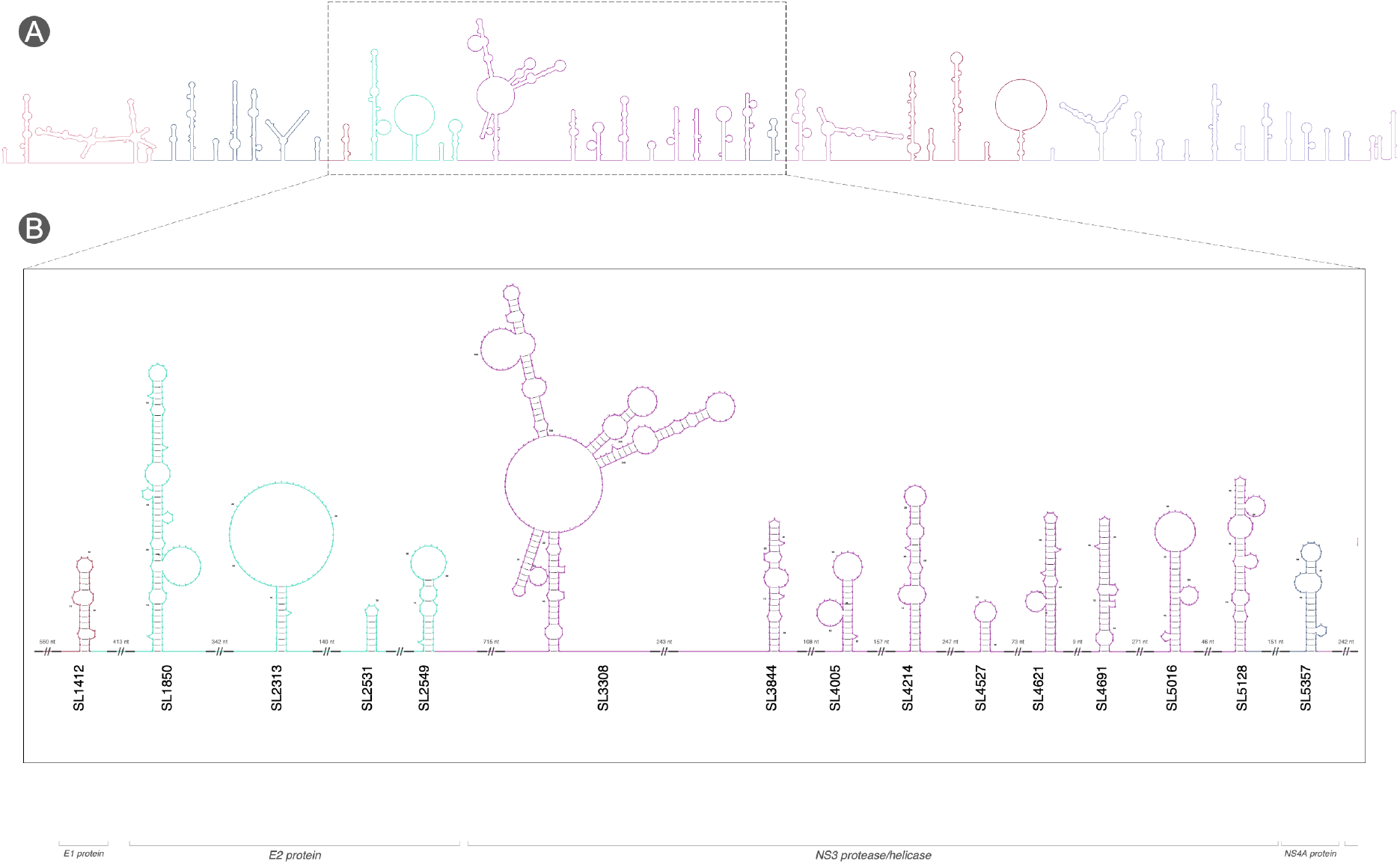
Genome-wide RNA structure annotation of Hepacivirus from a Stockholm alignment. **(A)** Overview of the full genome showing all RNA secondary structures as thumbnails along the genomic coordinate axis; a dashed box indicates the region expanded in panel B. **(B)** Zoomed-in view of the selected region. Each RNA structure is coloured according to its genomic region, with protein-coding regions labelled at the bottom of the panel (e.g., E1 protein, E2 protein, NS3 protease/helicase). Structure identifiers (e.g., SL1412, SL2313) are displayed beneath each diagram. Gaps between adjacent structures are indicated by double slashes (//) on the genomic axis, with the number of skipped nucleotides shown above (e.g., 560 nt, 413 nt).

`$r2dt.py stockholm examples/hepacivirus_hominis_alignment.stk output/hcv-stockholm/`

Overall, the use of multiple sequence alignments for RNA secondary structure prediction is a powerful strategy for uncovering evolutionarily conserved RNA architecture, particularly in viral genomes where sequence-based approaches alone are insufficient.

## 3 Programmatic access to Rfam

While the Rfam website provides a user-friendly means to visualise and interact with the data stored in the database, Rfam also provides a set of RESTful APIs which allow scripts or automated tools to access the data programmatically^4^. Rfam provides access to a comprehensive dataset of RNA Viruses in collaboration with the European Virus Bioinformatics Centre (EVBC). In the following examples we will look at the Coronavirus 3’ stem-loop II-like motif (s2m) family RF00164. This secondary structure motif also appears in the 3′ untranslated region (3′ UTR) of astrovirus and equine rhinovirus genomes and while its specific function remains unclear, 3′UTR structures in viruses are known to be involved in viral replication and genome packaging.

### 3.1 Retrieve family data with API

#### 3.1.1 Family description

Using the family endpoint, data for an individual Rfam family can be retrieved. The Rfam accession (in our example RF00164) or identifier (s2m) and the desired output format (HTML, JSON, or XML) are specified in the URL. The API can be accessed directly using curl:

$ curl “https://rfam.org/family/RF00164?content-type=application/json“

or using Python to retrieve information from an Rfam family:

~~~
import json
import requests
r = requests.get(‘https://rfam.org/family/RF00164?content-type=application/json’)
data = r.json()
# Pretty print the whole response
print(json.dumps(data, indent=2))
# Or just the accession
print(data[‘rfam’][‘acc’])
~~~

For brevity, most of the following examples will only demonstrate the curl method to return data, but these can easily be adapted to follow the same pattern as the Python script. The Family endpoint returns general information about an Rfam family, including a description, curation details, and search parameters.

#### 3.1.2 Accession and ID

The API offers additional endpoints that allow users to retrieve specific datasets relevant to their research. For example, use the “id” parameter to return the accession for family RF00164:

~~~
$ curl ‘https://rfam.org/family/RF00164/id‘
s2m
~~~

Or do the reverse by using the “acc” parameter to obtain the Rfam ID for the accession:

~~~
$ curl ‘https://rfam.org/family/RF00164/acc‘
RF00164
~~~

#### 3.1.3 Secondary structure images

The `/image/` endpoints return the schematic secondary structure images for the family in SVG format. Rfam provides several types of secondary structure diagrams to visualise different aspects of RNA structure and conservation. These include sequence conservation (cons), basepair conservation (fcbp), covariation analysis (cov), relative entropy (ent), maximum covariance model parse (maxcm), normal secondary structure (norm), R-scape [26] analysis of the Rfam SEED alignment (rscape), and secondary structure predicted by R-scape based on the Rfam SEED alignment (rscape-cacofold).

Rscape for RF03116 is visualised in Fig. 6 using a Jupyter notebook:

**Figure 6.**
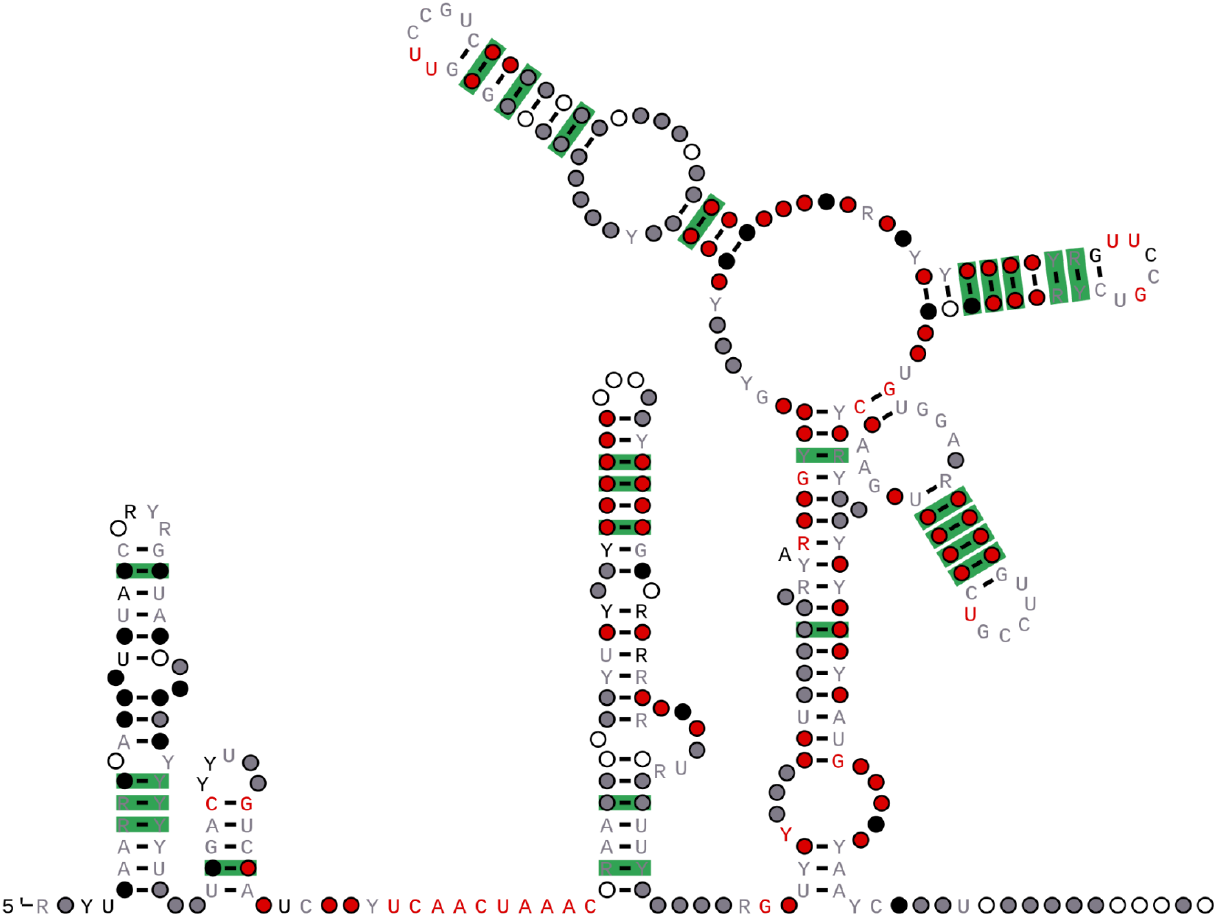
This Rfam family (RF03116) represents the secondary structure of the 5’-UTR of Alphacoronavirus. The secondary structure model highlights the statistically significant covarying base pairs in green. The R-scape algorithm is used to distinguish true structural covariance from the analysis of the SEED alignment [26]. Colours indicate basepair and nucleotide conservation: green, statistically significant basepair with covariation; red, 97% conserved nucleotide; black, 90% conserved nucleotide; grey, 75% conserved nucleotide; white, 50% conserved nucleotide. Nucleotide codes: R, A or G; Y, C or U.

~~~
from IPython.display import SVG, display
import requests
display(SVG(requests.get(‘https://rfam.org/family/RF03116/image/rscape’).content))
~~~

#### 3.1.4 Covariance models

The covariance model can be obtained with the `/cm` parameter:

~~~
$ curl ‘https://rfam.org/family/RF00164/cm‘
INFERNAL1/a [1.1.5 | Sep 2023]
NAME s2m
ACC RF00164
DESC Coronavirus 3’ stem-loop II-like motif (s2m)
STATES 133
NODES 33
CLEN 43
W 57
ALPH RNA
…
~~~

#### 3.1.5 Sequence regions

Rfam sequence regions are the specific coordinates in genomic sequences where the RNA family members have been identified. The data is available as a dataframe with columns for sequence accession, bit score, region start, region end, sequence description, species and NCBI tax ID and can be obtained with the /regions parameter as plain text or as xml with the addition of the `?content-type=text/xml` querystring:

$curl‘https://rfam.org/family/RF00164/regions?content-type=text%2Fxml‘

#### 3.1.6 Phylogenetic trees

The Rfam phylogenetic tree data charts the evolutionary relationship between members of the family based on the SEED alignment and built using the fasttree method. It is available in NHX (New Hampshire eXtended) format where the evolutionary distance between pairs of species is represented by branch length and a support value between 0 and 1. It can be obtained with the `/tree` parameter:

~~~
$ curl ‘https://rfam.org/family/RF00164/tree/‘
((_L06252.1/368-410_Infectious_bronchitis_v..[11120].4:0.02343,_AJ278335.1/370-
412_Infectious_bronchitis_v..[11120].6:0.00055)0.910:0.02313,(_AF111997.1/1443-
1485_Turkey_coronavirus[11152].2:0.02343,(_AJ619592.1/83-
125_Pheasant_coronavirus[258781].2:0.00055,_AY641576.1/27283-
27325_Infectious_bronchitis_v..[11120].3:0.02332)0.840:0.02332)0.510:0.00054,
…
~~~

A `.png` image of the phylogenetic tree can also be downloaded to a location specified with the `-–output` option. Choosing to use the `/species/` parameter will download an image with species names labelled, or choosing `/acc/` will download an image with species accessions labelled:

~~~
$ curl ‘https://rfam.org/family/RF00164/tree/label/species/image‘ --output
∼/RF00164_tree_label_species.png
~~~

#### 3.1.7 Structural mapping

The structural mapping data endpoint `/structures/` provides the Protein Data Bank (PDB) ids of 3D structural files of the RNA molecules that the family matches. Our example family RF00164 matches residues 4-46 of the A chain of the RNA structure with PDB ID of 1XJR. The data can be returned in either JSON format by appending the request with the ?content-type=application/json querystring or XML with the `?content-type=application/json` querystring. For example:

~~~
$ curl ‘https://rfam.org/family/RF00164/structures?content-type=application/json‘
{“mapping”:[{“rfam_acc”:”RF00164”,”bit_score”:65.6,”pdb_start”:4,”evalue_score”:”4.4e-
20”,”chain”:”A”,”pdb_id”:”1xjr”,”cm_start”:1,”pdb_end”:46,”cm_end”:43}]}
~~~

The PDB ID can then be used to view more information about the RNA structure on the Protein Data Bank website by appending it to the https://www.ebi.ac.uk/pdbe/entry/pdb/ url and visiting the webpage in a browser. In our example, at https://www.ebi.ac.uk/pdbe/entry/pdb/1xjr there is information available about publications, and biochemical data including complexes, ligands and macromolecules.

#### 3.1.8 Alignments

The SEED alignment of the family is available from the `/alignment/stockholm` endpoint in classic Stockholm format, or pfam Stockholm format (with sequences on a single line) from the `/alignment/pfam` endpoint. Alternatively the data can be obtained in ungapped or gapped FASTA format from the `/alignment/fasta` and `/alignment/fastau` endpoints respectively.

~~~
$ curl ‘https://rfam.org/family/RF00164/alignment/stockholm‘
# STOCKHOLM 1.0
#=GF ID s2m
#=GF AC RF00164
#=GF DE Coronavirus 3’ stem-loop II-like motif (s2m)
#=GF AU Nawrocki E; 0000-0002-2497-3427
#=GF GA 50.0
#=GF NC 47.7
#=GF TC 51.6
#=GF SE PMID:9568965; Griffiths-Jones SR, Moxon SJ
#=GF SS Published; PMID:15630477
…
~~~

As with the phylogenetic tree images, the alignment files can be downloaded with addition of the `–output` option:

$ curl ‘https://rfam.org/family/RF00164/alignment/stockholm‘ --output ∼/RF00164.sto

Appending the url with the `?gzip=1` querystring will download a compressed version of the file:

$ curl ‘https://rfam.org/family/RF00164/alignment/stockholm?gzip=1‘ --output ∼/RF00164.sto.gz

The Rfam website features an alignment viewer which parses the same data to provide a visualisation of the SEED alignment (**Fig. 7**).

**Figure 7.**
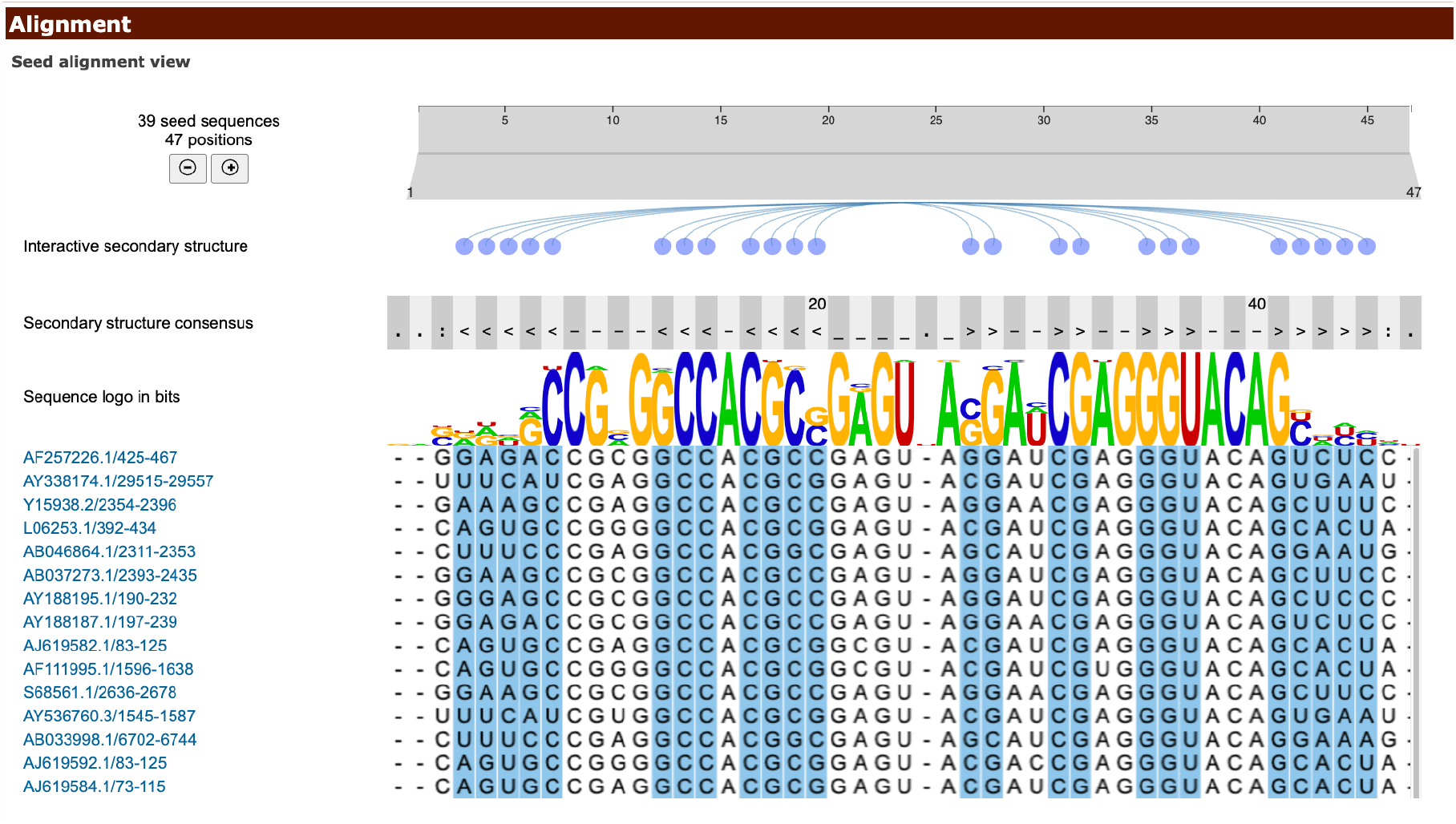
SEED alignment view of Rfam family RF00164 implemented with the Nightingale framework [27], available at https://rfam.org/family/RF00164#tabview=tab2.

### 3.2 Search your sequence using the API

Querying a nucleotide sequence against the Rfam covariance model library identifies regions in your sequence that belong to one of our classified RNA families^5^.

As an example, we are going to select a 281-nucleotide Pestivirus giraffe-1 H138 internal ribosome entry site (IRES) sequence from RNAcentral [28] https://rnacentral.org/rna/URS0000D7A5FF/119222, and we save the sequence in a FASTA formatted file.

~~~
$ curl -X ‘POST’ ‘https://batch.rfam.org/submit-job‘ -H ‘accept: application/json’ -F
‘sequence_file=@URS0000D7A5FF_119222.fa’
{“resultURL”:”https://batch.rfam.org/result/infernal_cmscan-R20260218-125553-0949-88811294-p1m“,”jobId”:”infernal_cmscan-R20260218-125553-0949-88811294-p1m”}
~~~

We can programmatically get the full output by:

$ curl -s ‘https://batch.rfam.org/result/infernal_cmscan-R20260218-125553-0949-88811294-p1m‘ | jq ‘.’

We can obtain details for each hit, in a table format, by:

~~~
$ curl -s ‘https://batch.rfam.org/result/infernal_cmscan-R20260218-125553-0949-88811294-p1m‘ | jq -r ‘ [“Family”, “Accession”, “Score”, “E-value”, “Start”, “End”], (.hits | to_entries[] | [.key,
.value[0].acc, .value[0].score, .value[0].E, .value[0].start, .value[0].end]) | @tsv ‘ | column -t
Family Accession Score E-value Start End
IRES_Pesti RF00209 255.3 6.4E-76 1 273
~~~

As expected, the most significant hit corresponds to the Pestivirus internal ribosome entry site (IRES) Rfam family RF00209. Search results will be available for seven days.

### 3.3 Text search using the API

The Rfam text search enables exploration of the database by category, sorting results, and filtering by various facets such as RNA type, taxonomic distribution, or presence of 3D structures^6^. This search is powered by the EBI Search service, which provides a REST API for programmatic access [29].

#### 3.3.1 Searching for viral RNA families

To search for all coronavirus-related families and retrieve their descriptions along with the number of SEED and FULL sequences:

~~~
$ curl -s
‘https://www.ebi.ac.uk/ebisearch/ws/rest/rfam?query=coronavirus&format=json&fields=description,num_see
d,num_full’ | jq ‘.entries[] | {id, description: .fields.description[0], num_seed:
.fields.num_seed[0], num_full: .fields.num_full[0]}’
{
 “id”: “RF00507”,
 “description”: “Coronavirus frameshifting stimulation element”,
 “num_seed”: “51”,
 “num_full”: “76”
}
{
 “id”: “RF03116”,
 “description”: “Alphacoronavirus 5’UTR”,
 “num_seed”: “17”,
 “num_full”: “31”
}
…
~~~

#### 3.3.2 Retrieving viral families using Python

One can use Python to retrieve and tabulate Hepatitis C virus families:

~~~
import requests
url = ‘https://www.ebi.ac.uk/ebisearch/ws/rest/rfam’
params = {
 ‘query’: ‘hepatitis C virus’,
 ‘format’: ‘json’,
 ‘fields’: ‘description,num_seed,num_full,rna_type’
}
r = requests.get(url, params=params)
data = r.json()
print(f”Found {data[‘hitCount’]} families:”)
for entry in data[‘entries’]:
 print(f”{entry[‘id’]}: {entry[‘fields’][‘description’][0]}”)
~~~

#### 3.3.3 Filtering for viral cis-regulatory elements

To retrieve cis-regulatory RNA families found in viruses using an RNA type filter:

~~~
$ curl -s -G -H “Accept: application/json” --data-urlencode ‘query=rna_type:Cis-reg AND
tax_string:Viruses’ --data-urlencode ‘fields=description’ https://www.ebi.ac.uk/ebisearch/ws/rest/rfam
| jq ‘.hitCount’
3300
~~~

A full list of searchable fields is available at https://www.ebi.ac.uk/ebisearch/metadata.ebi?db=rfam.

## Availability

Rfam is freely available at https://rfam.org. The data can be accessed via an API, a public MySQL database and the FTP archive. An Rfam SVN repository with unreleased updates and newly-added families is available at svn.rfam.org. All code is available on GitHub under the Apache 2.0 licence at https://github.com/Rfam and https://github.com/R2DT-bio/R2DT. The code and materials for this chapter can be found at https://github.com/Rfam/rfam-viral-rna-structure.

## Acknowledgments

This work was supported by grants from the Biotechnology and Biological Sciences Research Council (BBSRC) [UKRI746:24BBR] and the Wellcome Trust [310300/Z/24/Z], and in part by the Intramural Research Program of the National Institutes of Health (NIH). The contributions of the NIH author are considered Works of the United States Government. The findings and conclusions presented in this paper are those of the author(s) and do not necessarily reflect the views of the NIH or the U.S. Department of Health and Human Services.

## Footnotes

1 R2DT is available at https://r2dt.bio/ and provides automated generation of secondary structure diagrams for RNA sequences. For viral genome examples, using a custom CM library, customising the stitched output, or integrating with other R2DT workflows, see https://docs.r2dt.bio/en/latest/viral-genomes.html.

2 Filtering out lab strains when collecting sequences for viruses is currently not provided within the NCBI Datasets CLI. However, it is possible to filter and download viral genomes directly from the NCBI Virus Database available at https://www.ncbi.nlm.nih.gov/labs/virus/vssi/#/.

3 Viral RNA families in Rfam are available at https://rfam.org/viruses. This page provides a comprehensive list of RNA families identified in viral genomes, organised by virus taxonomy. The Rfam team is currently working on adding RNA families from other viruses, such as *Filoviridae* (e.g. Ebolavirus) and *Rhabdoviridae* (e.g. Rabies viruses).

4 The Rfam documentation is available at https://docs.rfam.org/en/latest/, and provides comprehensive guides on using the database, understanding the data structure, and accessing various features. API-specific documentation can be found at https://docs.rfam.org/en/latest/api.html#data-access.

5 The Sequence Search service to find Rfam families within your sequence of interest is available at https://rfam.org/search#tabview=tab1. The results include secondary structures and significant matches to sequences in RNAcentral [28].

6 Documentation about the Rfam Text Search API can be found at https://docs.rfam.org/en/latest/searching-rfam.html#text-search-api. This API endpoint allows programmatic text-based searches across Rfam entries using keywords, enabling users to query family names, descriptions, literature references, and other metadata fields.

